# Competitive advantages of T-even phage lysis inhibition in response to secondary infection

**DOI:** 10.1101/2024.02.07.579269

**Authors:** Ulrik Hvid, Namiko Mitarai

## Abstract

T-even bacteriophages are known to employ lysis inhibition (LIN), where the lysis of an infected host is delayed in response to secondary adsorptions. Upon the eventual burst of the host, significantly more phage progenies are released. Here, we analysed the competitive advantage of LIN using a mathematical model. In batch culture, LIN provides a bigger phage yield at the end of the growth where all the hosts are infected due to an exceeding number of phage particles and, in addition, gives a competitive advantage against LIN mutants with rapid lysis by letting them adsorb to already infected hosts in the LIN state. By simulating plaque formation in a spatially structured environment, we show that, while LIN phages will produce a smaller zone of clearance, the area over which the phages spread is actually comparable to those without LIN. The analysis suggests that LIN induced by secondary adsorption is favourable in terms of competition, both in spatially homogeneous and inhomogeneous environments.

**Author Summary:** T-even bacteriophages can delay the lysis of their hosts when they detect more phages are adsorbing to the hosts, increasing the progeny production per host. Using a mathematical model, we provide a quantitative analysis of this strategy’s competitive advantages and disadvantages in different environments. The model predicts that phage adsorption to lysis-inhibited cells provides a significant competitive advantage to lysis-inhibiting phage against phages that quickly lyse the cells. We also find that secondary infection-triggered delay does not hinder the spreading of the phage in a lawn of uninfected cells, even though the apparent plaque size is small. The analysis suggests that lysis inhibition provides a robust competitive advantage for a virulent phage.

## Introduction

T-even phages such as T2, T4, and T6 that infect *Escherichia Coli* have long been used as a laboratory model system to study bacteriophages. They are well-studied, obligate lytic phages, i.e. an infected cell will invariably lyse to produce many progenies. A common feature that T-even phages exhibit is lysis inhibition (LIN) [1, 2]. LIN is a mechanism of latent-period extension and burst-size increase induced in a cell primarily infected by a T-even phage when the cell is superinfected by the same phage [2].

One of the characteristics of T-even phages is the small size of their plaques, i.e., the circular zone of cell destruction caused by a phage spreading in a growing bacterial lawn. Mutants of these phages that show significantly larger plaque morphology have been identified, and these mutants turned out to be deficient in LIN. Hence, they are called *r*-mutants for ‘rapid lysis’ [1]. The study of *r*-mutants has contributed significantly to today’s understanding of LIN and their molecular mechanisms [3, 4].

LIN is an interesting phenotype since the phage responds to secondary adsorptions, which signal that there are more phage particles than available hosts in the local environment. LIN is considered a strategy to switch behaviour when the environment changes from an excess host situation to the opposite, a phage excess environment. In the former, exponential spread among new hosts through short latency time and multiple rounds of burst is beneficial if the burst size is a constant. In reality, there is a trade-off between the latency time and the burst size; for *λ* phage, it was reported that the burst size increases linearly with the latency time, and an optimal latency time was found to maximize the fitness under this trade-off [5]. When there are no more uninfected hosts left, on the other hand, it is better to milk the last host to produce as many phage particles as possible per infected host. Because there is a trade-off between the lysis time and the burst size [5–8], switching based on the phage/bacteria ratio becomes a beneficial strategy.

We can ask further if LIN has any more advantages or disadvantages. One of the obvious phenotypes of LIN is their small plaque size, which distinguishes them from the *r*-mutant. This could suggest that LIN is delaying the spreading of phage in a structured environment because the phage concentration becomes locally high, even if there are abundant hosts further away. If so, it is a disadvantage of LIN. Interestingly, however, it has been previously suggested by Abedon [9] that the infection front of a T4-plaque spreads a comparable distance to that of an *r*-mutant, but the outer zone stays turbid because the cells are largely in LIN state and not lysed. The same author also suggested that LIN may provide an additional competitive advantage by adsorbing and deactivating competitor phages while in the LIN state.

In this paper, we study LIN’s competitive advantages and disadvantages by using mathematical models. The model allows us to control various features of LIN, which helps test the effect of each feature. We compare the behaviours of the phages capable of LIN (we call them LIN-phages in the following) and the *r*-mutants and show that the inactivation of competitor phages by adsorption in LIN state can be indeed a significant benefit during competition in a well-mixed culture with high bacterial concentration. We also show that the mathematical model indeed allows for the spread of infection front quite far compared to the zone of clearance for secondary adsorption-mediated LIN phages because the diffusion of the phage ensures the existence of a low secondary adsorption zone at the front of the expanding plaque.

## Models

We first present a mathematical model for *r*-mutant or LIN-phage spreading among suceptable host bacteria to simulate a well-mixed batch culture. We then extend the model to include spatial structure to simulate the plaque formation [10].

### M0 - Model for *r*-mutant

We present a simple model without LIN, which is a model for *r*-mutants. This model will be referred to as M0. We consider the concentration of the uninfected bacteria *B*, free phage particles *R*, and the infected bacteria 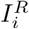, where the subscript *i* denotes the progress in the lysis process. We also assume that the bacteria and phage growth are affected by the concentration of the nutrient *n*, where the growth rate at a given *n, g*(*n*), is given by Monod’s growth law [11] as

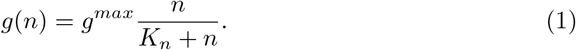

Here, *g*^*max*^ is the maximum growth rate when the nutrient availability is unlimited, and *K*_*n*_ is the Monod constant that characterizes the nutrient concentration at the half-maximum growth rate.

We assume Lotka-Volterra type interaction with an adsorption rate *η* for phage infection term. We assume both uninfected and infected cells adsorb and inactivate free phages. An infected suceptable bacterium will burst *β*(*n*) new phage particles after the latency time *τ*(*n*), which are nutrient concentration dependent. In order to have the latency time distribution centred around *τ*(*n*), the infected bacteria go through *N* sequential steps before the cell burst [10]. These assumptions are summarized into the following model equations and visualized in Fig. 1:

**Fig. 1:**
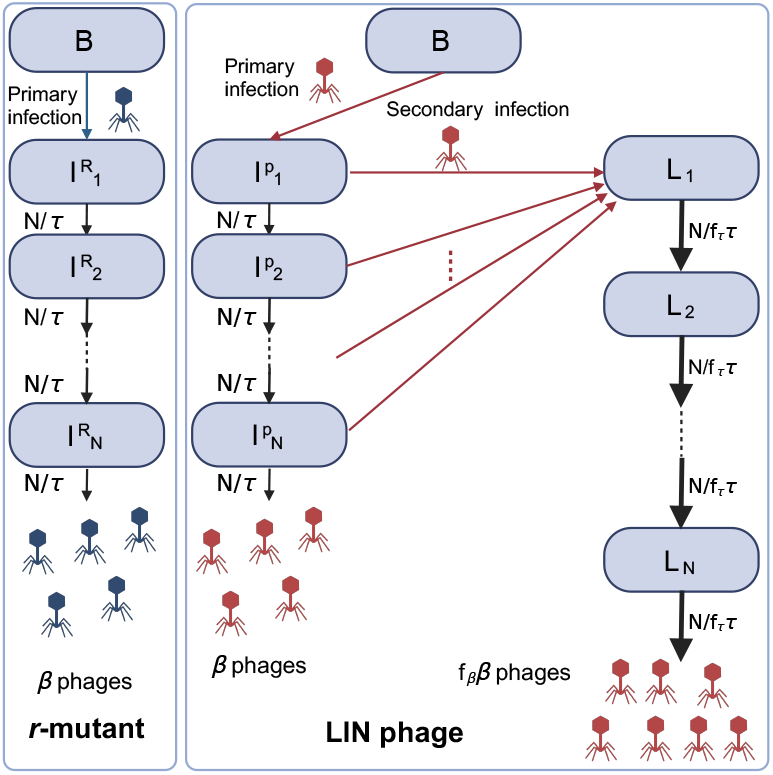
Diagram to illustrate state transitions in the model. Created with BioRender.com.

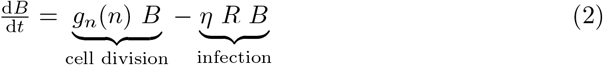

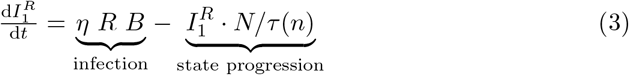

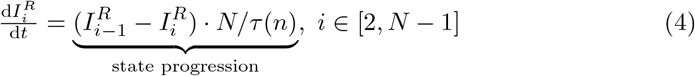

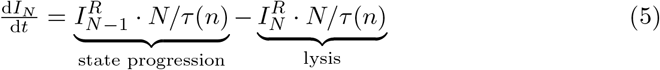

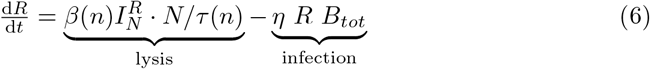

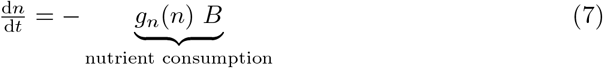

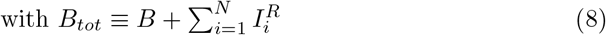

Note that the nutrient consumption rate is taken to be equal to the bacterial growth rate (Compare the first term in the right-hand side of eqs. (2)) and (7), i.e., the unit concentration of nutrient is taken to be the nutrient needed to produce one unit concentration of bacterium.

Inspired by [7, 12], we assume the functional form of the nutrient dependence of the phage parameters as follows:

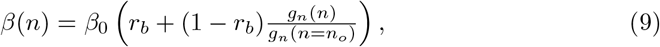

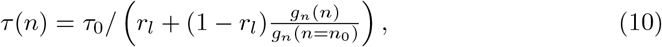

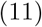

Here *r*_*b*_ and *r*_*l*_ are the ratios of maximal to minimal burst size and latency time, respectively, and *n*_0_ is the initial nutrient concentration.

### M1 - simple model of LIN

Next, we introduce a simple model that includes LIN. This model is for LIN-phage and will be referred to as M1. When a LIN-phage (concentration *P*) infects an uninfected bacterium (*B*), the cell will transition to the infected state 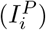, and, in the absence of further phage adsorption, the cell will proceed towards lysis as in M0. When an infected cell 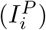 is superinfected for the first time, the cell transition to the first step of the lysis-inhibited state (*L*_1_). For simplicity, we do not take into the effect of further phage adsorption. Instead, similar to the infected cell in model M0, a lysis-inhibited cell will go through *N* intermediate step (denoted by *L*_*i*_), with a total latency time *τ*_*I,l*_ = *f*_*τ*_ *τ*_*l*_. The parameter *f*_*τ*_ characterises how long the additional latency time is compared to *r*-mutant’s latency time *τ*_*l*_, and we focus on the case *f*_*τ*_ ≥ 1. Note that this ignores the possible dependence of the latency time on the number and timing of additional phage adsorptions [3]. When the cell in LIN-state lyses, it releases *β*_*I*_ = *f*_*β*_*β* phages, where *f*_*β*_(≥ 1) describes the fold-increase of the burst size after LIN. The state transitions are visualized in Fig. 1. In reality, a positive correlation between the increase of the latency time and the increase of the burst size has been observed [5, 13]. However, in the present analysis, we control the two parameters independently to allow the model to explore the pros and cons of LIN fully. We assume that the adsorption rate of LIN-phage *P* is the same as that of *R*. Importantly, bacteria in a LIN-state keep adsorbing phages, but that will not change the state of the cell any further. This means that the phages absorbed by LIN-state bacteria are simply inactivated without further production of progeny and purely wasted, similar to the phage adsorption to bacteria debris that has been proposed to play a role in phage-bacteria coexistence [14].

This results in the following modification of equations:

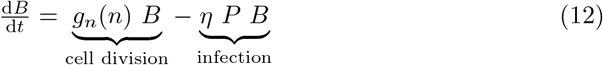

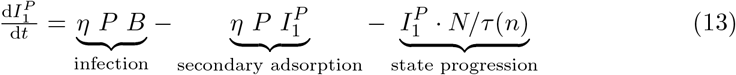

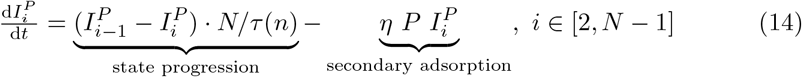

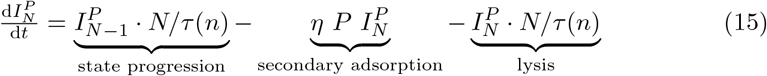

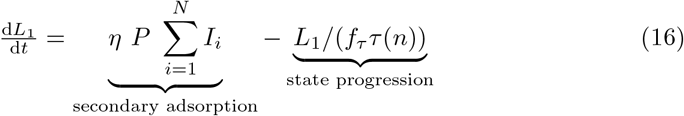

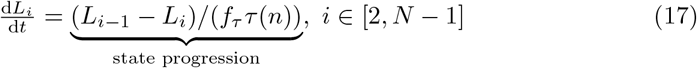

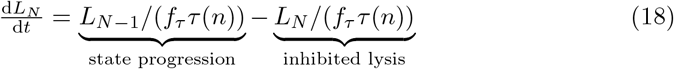

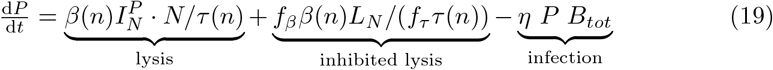

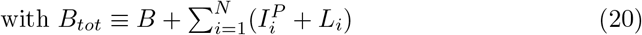

The equation for the nutrient concentration, *n*, remains the same as (7).

### MC1 - Model for competition between LIN-phage and *r*-mutant

Finally, we present a model where both the LIN-phage (concentration *P*) and the *r*-mutant (*R*) compete in the same culture. We call this model MC1. We assume that the first infection determines which phage strain will eventually be produced upon burst. This is because of the superinfection exclusion, whereby the T4 genes *imm* and *sp*, expressed by the primarily infecting phage, block the genome of a secondarily infecting phage from entering the cytoplasm [9]. Furthermore, it has been proposed that the entrance of DNA to periplasm is the signal to cause LIN, possibly by stabilizing the holin-antiholin complexes [15, 16]. Therefore, we assume that if a cell is first infected by LIN-phage *P* (i.e. in 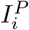 state), secondary adsorption by any phage (*P* or *R*) will bring the cell to LIN-state in the main text (We study the case where only *P* can trigger LIN in the supplementary material Fig. S1). Conversely, if the first infection is by an *r*-mutant *R*, then secondary adsorption does not cause LIN, regardless of the superinfecting strain. The resulting set of equations is a straightforward extension of M0 and M1 as follows:

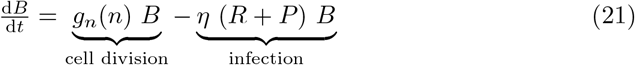

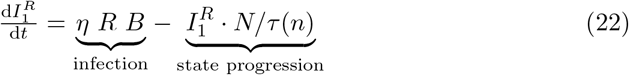

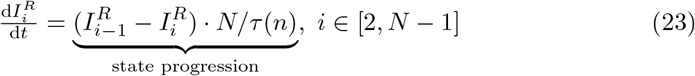

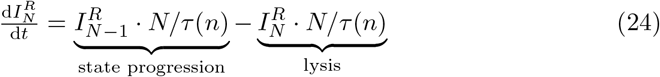

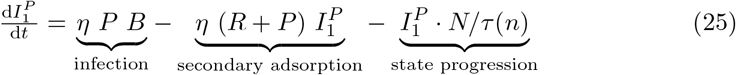

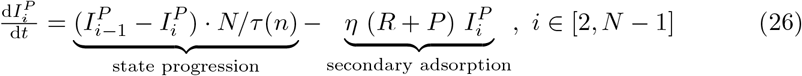

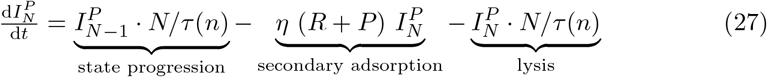

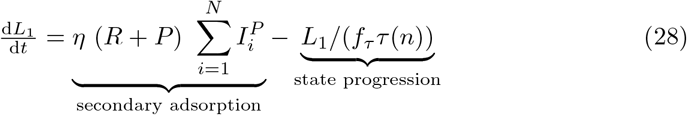

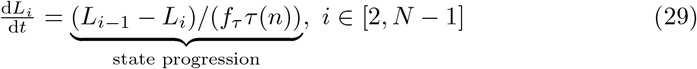

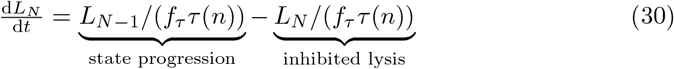

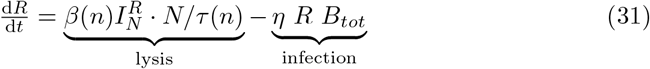

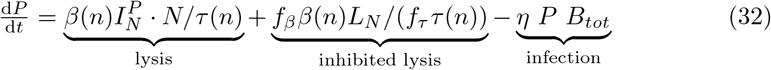

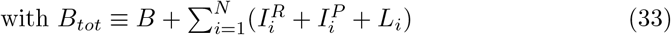

### Model for plaque formation

We model a plaque formation experiment setup, where a large number of bacteria and a small number of phages are mixed in a soft agar and cast in a thin layer on a hard agar plate containing the nutrient. The subsequent overnight incubation results in the growth of bacteria where there is no phage, while local growth and diffusive spreading of phage create a zone of cell destruction that appears as a plaque [17]. Bacteria cannot swim visibly in the high viscosity of soft agar [18]. Hence, each initially cast bacterium divides locally to form a microcolony [19].

Here, we modify the plaque formation model proposed [10] to include the LIN. For simplicity, the model assumes a locally well-mixed population and does not consider the local growth of bacteria into microcolonies that may provide extra protection against phage attack [19–21]. We use the same symbols as the well-mixed models M0 and M1 to denote the different populations, but now they are expressed in the unit of number per area by integrating the concentration over the thin soft-agar of thickness Δ*a*. We call the spatial version of M0 as MP0, and the spatial version of M1 as MP1. Here, the variables are functions of both time *t* and the position 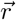. Diffusion terms are added for the nutrients and phages. The resulting model equations are parallel to the ones in M0 for *r*-mutant or M1 for LIN-phage, but the total derivative by time in the left-hand side of equations should be replaced with partial derivative by time, and the diffusion terms should be added to the right-hand side of Eqs. (6), (7), and (19), with diffusion constant for phage *D*_*p*_ (assumed to be the same for both *r*-muant and LIN-phage) and that for nutrient *D*_*n*_. Furthermore, the phage adsorption rate *η* needs to be divided by Δ*a*, reflecting that the population is integrated over this depth. The full equations are explicitly presented in the supplementary text SI.

Finally, to reflect the change of unit of the nutrient per volume to per area, the symbol for Monod constant is changed from *K*_*n*_ in the well-mixed Eq. (1) to *k*_*n*_ for the spatial model (used in the parameters in Table 1 and 2), respectively. To stop the plaque spreading when the nutrient runs out, we set the latency time to diverge when no nutrient is available by setting *r*_*l*_ = 0 in eq. (10) [10].

**Table 1:** Default parameter values and initial values for the simulation of well-mixed batch culture (M0, M1, MC1). By default, the initial value of infected bacteria is zero.

| Parameter | Description | Default value | Source |
| --- | --- | --- | --- |
| $g_n^{max}$ | Max growth rate of bacteria | $0.034 \text{ min}^{-1}$ | 20 min doubling time |
| $K_n$ | Nutrients for half-max growth rate | $n_0/5$ | [10] |
| $\eta$ | Adsorption rate | $5 \times 10^{-10} \text{ ml/min}$ | [22], [23] |
| $\tau_0$ | Minimum lysis-time pr. substate | $(20 \text{ min})/N$ | [7], [22] |
| $f_\tau$ | LIN latency time/(latency time w/o LIN) | varied | Only in M1 and MC1 |
| $\beta_0$ | Maximum burst size | 150 | [22] |
| $f_\beta$ | LIN burst size/(burst size w/o LIN) | varied | Only in M1 and MC1 |
| $r_l$ | Ratio of maximum to minimum latency time | 0.5 | [7] |
| $r_b$ | Ratio of maximum to minimum burst size | 0.1 | [7] |
| $N$ | Number of infected states in model | 10 | [12] |
| $B_0$ | Initial concentration of bacteria | $10^6 \text{ ml}^{-1}$ | Free parameter |
| $P_0$ (or $R_0$) | Initial concentration of phages | $10^3 \text{ ml}^{-1}$ | Free parameter |
| $n_0$ | Initial concentration of nutrients | $10^9 \text{ ml}^{-1}$ | Yields $10^9 \text{ ml}^{-1}$ cells |

**Table 2:** Default parameter values and initial values for additional/changed parameters/values in the plaque formation simulation (MP0, MP1.)

| Parameter | Description | Default value | Source |
| --- | --- | --- | --- |
| $\Delta a$ | Soft agar thickness | 0.5mm | [10] |
| $r_l$ | Ratio of maximum to minimum latency time | 0 | To stop plaque development [10] |
| $D_p$ | phage diffusion coefficient | $240\mu\text{m}^2/\text{min}$ | In agarose gel [24] |
| $D_n$ | nutrient diffusion coefficient | $5 \times 10^4 \mu\text{m}^2/\text{min}$ | [12] |
| $k_n$ | Nutrients for half-max growth rate | $n_0/5$ | [10] |
| $B_0$ | Initial concentration of bacteria | $1/400/\mu\text{m}^2$ | Free parameter |
| $n_0$ | Initial concentration of nutrients | $0.5/\mu\text{m}^2$ | Free parameter |

We focus on a single plaque simulation and set the coordinate’s origin at the plaque’s centre. We assume circular symmetry of the plaque structure, which allows us to solve the problem in polar coordinates and focus on the variation along the radial direction, *r*. At a distant outer boundary, *r* = *R*_*max*_, we impose reflecting boundary conditions 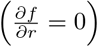 for both phage and nutrient density.

### Parameters

We here list the parameter values used in the simulations.

Initially, bacteria and nutrients are uniformly distributed [*B*(*r*, 0) = *B*0 and *n*(*r*, 0) = *n*0], and one freshly infected bacterium is placed in the middle 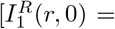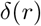 or 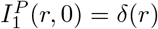 with the Dirac delta function *δ*(*x*)], and there are no free phage [*P*(*r*, 0) = 0].

### Numerical integration

The integration of well-mixed models was done by *solve ivp* under scipy.integrate with fourth-order Runge-Kutta method.

## Results

### Burst size increase is necessary for LIN-phage to have an advantage over *r*-mutant in separate culture growth

The simulation of the phage and bacteria growth in well-mixed batch culture using the model described in the model section and summarized in Fig. 1 is shown in Fig. 2 for *r*-mutant (A) and LIN-phage (B). Here, the LIN parameters are set to *f*_*τ*_ = *f*_*β*_ = 5, i.e., a secondary adsorption results in about a 5-fold increase in latency time as well as burst size. Naturally, the two cases show quantitatively similar trajectories until free phage concentration becomes comparable with uninfected bacteria (around 2 hours with the current setup), where secondary adsorptions become frequent. After that, the free phage increase slows down in the LIN-phage case due to the increasing number of cells entering the LIN-state rather than immediately lysing (Fig. 2B around 2.5 hours).

**Fig. 2:**
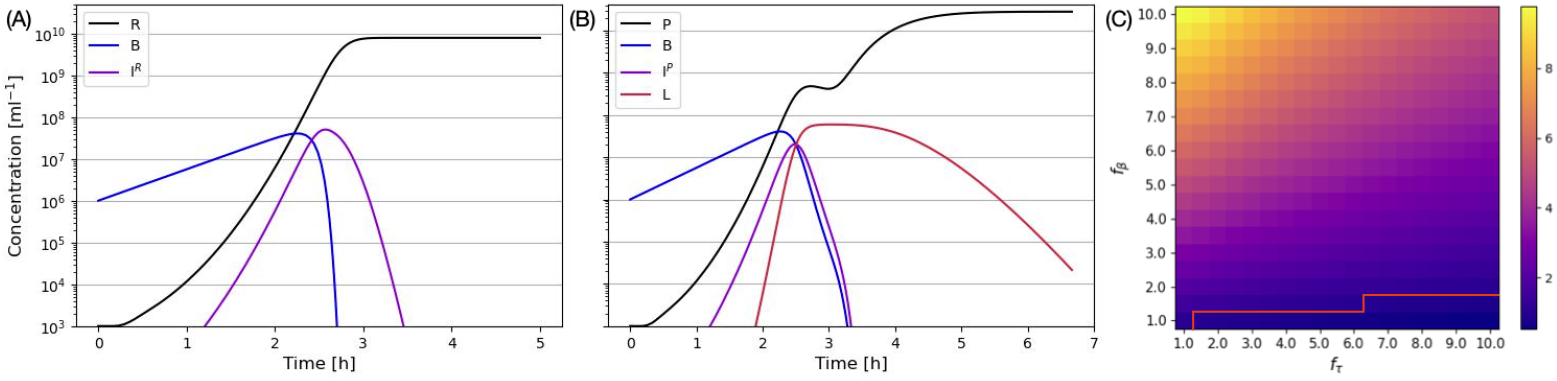
Well-mixed culture with and without LIN. Left top (A): Without lysis inhibition (M0). 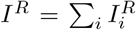 shows the sum of all infected but not lysed cells. (B): With lysis inhibition (M1) with 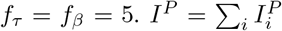 shows the sum of all cells infected once but not lysed, and *L* = Σ_*i*_ *L*_*i*_ shows the sum of lysis-inhibited cells. (C): The final phage ratio between LIN-phage (*P*) and r-mutant (*R*) after 20 hours. The red line shows the contour of the ratio being one. LIN-phage is dominating in most of the parameter space.

In the current parameter set, those bacteria start to lyse around 4 hours. Since we assume a higher burst size for LIN-cells, LIN-phages reach a higher concentration of free phages than *r*-mutant at the end of the simulation. This final advantage is due to the trade-off between the latency time and burst size assumed in the parameter choice of *f*_*τ*_ = *f*_*β*_ > 1. There is, however, a downside of the LIN due to phage adsorption by bacteria in LIN-state: If *f*_*τ*_ is very long, there is a clear drop of the free phage concentration before the burst of LIN cells, depicted as a small decrease of free phage around 2.5 hours in Fig. 2B.

In order to test this trade-off in LIN, we tested the advantage of LIN-phages by varying *f*_*τ*_ and *f*_*β*_ independently. Figure 2C shows the ratio of the final number of free phages against that of *r*-mutant phage after 20 hours when most of the infected cells were lysed. We clearly see that LIN phage has an advantage against *r*-mutant (i.e., the ratio is bigger than one) only if *f*_*β*_ > 1, and the bigger *f*_*τ*_, the bigger *f*_*β*_ is needed to have the advantage.

### LIN provides a competitive advantage even without burst size increase in a mixed culture competition

We next simulated the growth where the LIN-phages and *r*-mutants are mixed in the same culture and studied the outcome of the direct competition. Here, there are two critical assumptions in the model. One is that the primary infection determines which phages will eventually be produced. This assumption is because of superinfection exclusion, where the expression of *imm* and *sp* genes block the genome of a secondarily infecting phage from entering the cytoplasm [9], and thus from expression. Another assumption is that the infected cells can still adsorb free phages independent of the phage types (the genome will stay in periplasm), and secondary adsorption of any phage can trigger LIN if and only if the cell is infected by LIN-phage first.

Figure 3A shows the competition of the phages with the parameters presented in the independent culture simulation Fig. 2AB. While most of the behaviours appear as a simple overlay of the independent culture infections, a clear difference is seen in the concentration of free *r*-mutant phages: It peaks around 2.5 hours and then somewhat decreases. Note that the plot is the log scale, and the decrease is almost 50%. This is because the cells in LIN-state keep absorbing the phages for a long time. Therefore, LIN gives an additional competitive advantage if the two types of phages are competing in the same environment, as suggested in [9].

**Fig. 3:**
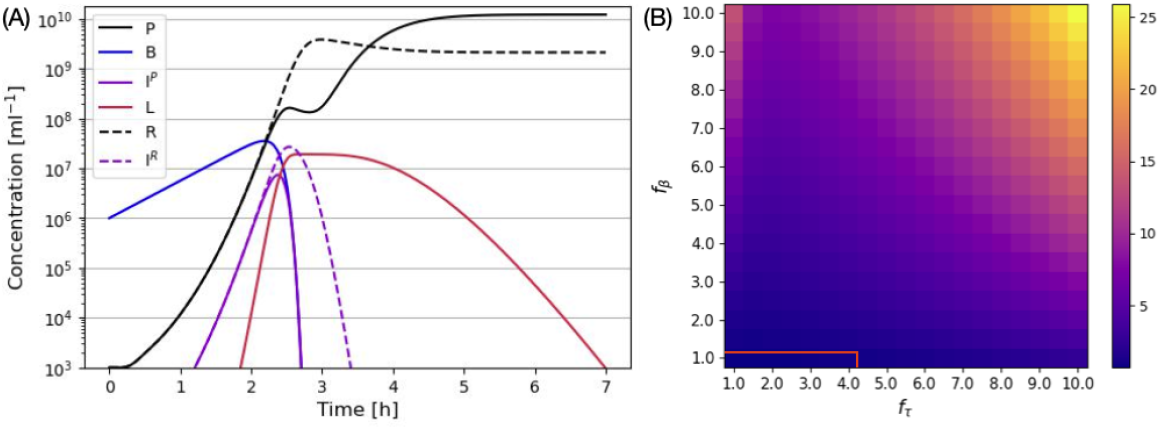
Competition of r-mutant and LIN-phage in a mixed culture. (A) An example time course of competition between *r*-mutant and LIN-phage with *f*_*τ*_ = *f*_*β*_ = 5 in a mixed culture. (B) The final phage ratio between r-mutant (*R*) and LIN-phage (*P*) after 20 hours in a mixed culture. The red line shows the contour of the ratio being one.

In Fig. 3B, we show the ratio between the free LIN phage and the rapid lysis phage *P/R* after 20 hours (when most of the infected cells were lysed) for various values of *f*_*r*_ and *f*_*τ*_. Except for rather short lysis time and no burst size increase combination (below *P/R* = 1 contour), we see that the LIN phages dominate the population in the end. It is especially worth noting that, as long as the LIN latency time is significantly longer than the *r*-mutant latency time, the LIN-phage wins over the *r*-mutant even if the burst size does not increase (*f*_*β*_ = 1).

While this result is based on the model that assumes the secondary adsorption of *r*-mutant can also trigger LIN, it is possible that some *r*-mutant cannot trigger LIN; for example, secondary adsorption of T2 *r*-mutant by a cell primary infected by T4 phage gave little LIN [2]. Therefore, we also tested the model where the secondary adsorption of *r*-mutant does not trigger LIN, and the result is shown in the supplementary Fig. S1. We observed an even stronger competitive advantage of LIN-phage against *r*-mutant because this ensures that LIN happens only after the concentration of LIN-phage becomes high enough; hence, LIN-phage does not fall behind to claim the free bacteria in comparison to *r*-mutant. This indicates that the competitive advantage of LIN by inactivating the competitor phage is robust against which phage can trigger LIN.

### LIN-phage plaque expands as far as *r*-mutant phage plaque

Next, we analyse the effect of LIN on plaque expansion. We extend the ordinary differential equation model for batch culture growth to partial differential equations to include the spatial structure, where the diffusion of the phages and nutrients is considered (Models and Methods for detail). In order for the plaque expansion to come to a halt when the nutrient runs out, we set *r*_*l*_ = 0, which results in divergent lysis time with decreasing nutrients.

Figure 4 shows the time series of plaque expansion dynamics for *r*-mutant (A) and LIN-phage (B). The horizontal axis shows the distance from the plaque centre, where an infected cell is placed at time zero. The phages produced upon burst diffuse and infect new cells, expanding the infected and the lysed zone over time. The area where the host cells are lysed determines the final visible plaque size. Importantly, for a LIN-phage plaque, when cells are in LIN-state, they will appear as a part of the lawn, though careful observation may reveal that the edges of the plaque are translucent [9]. The phage profile for LIN phage shows an additional peak for a later time (See Fig. 4B after 180min). The peak closer to the plaque center is due to the delayed lysis with a bigger burst size of some of the LIN-ed cells, while the smaller peak closer to the plaque edge is due to the cells bursting without delay. To see how this lawn of cells extends, we plotted the total concentration of uninfected and infected bacteria *B*_*tot*_ as the dotted lines in Fig. 4AB. We see that the LIN-phage (B) has a smaller radius of cell lysis than that of *r*-mutant (A), i.e., smaller plaque. The change of the total bacteria concentration over distance is more gradual in the LIN-phage, likely to give a translucent edge around the zone of killing. Interestingly, however, the front of uninfected bacteria in LIN-phage plaque and *r*-mutant plaques give similar profiles.

**Fig. 4:**
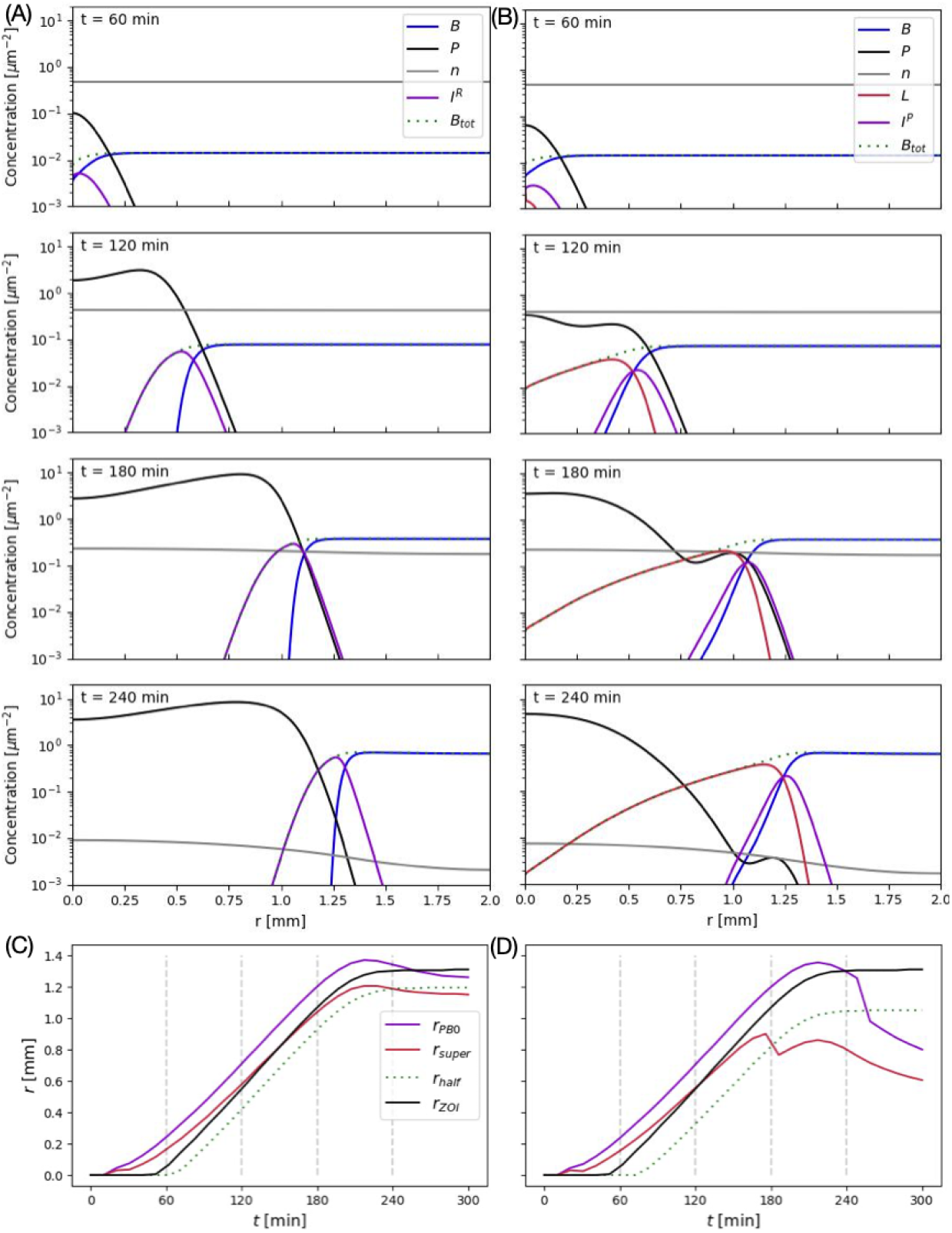
Plaque expansion of *r*-mutant phage and LIN-phage. (A) *r*-mutant plaque. (B) LIN-phage plaque with *f*_*τ*_ = *f*_*β*_ = 5. The dotted line shows the total concentration of bacteria cells (which is assumed to correspond to the visible bacterial lawn in the plate). (C) and (D): The time evolution *r*_*PB*0_, *r*_*super*_, *r*_*half*_, and *r*_*ZOI*_. (C) for *r*-mutant, and (D) for LIN-phage with *f*_*τ*_ = *f*_*β*_ = 5. Vertical dotted lines mark the time points corresponding to the plots in (A) and (B).

In other words, in the simulation, the LIN-phage managed to infect almost as large an area as the *r*-mutant phage, but the cells are not lysed and the plaque size appears to be small.

The expansion of the phage-infected area for LIN-phages is the natural consequence of the secondary adsorption-induced LIN. Since the released phages diffuse over space before infection, the free phage concentration decreases with the distance from the last burst site. Consequently, the phage concentration is high close to the burst site, giving LIN, while phage concentration drops to one phage per bacteria deeper into the bacterial lawn, ensuring that some cells in the infection front are not lysis-inhibited. Hence, the cells in the front of the infection keep bursting with short latency time, ensuring the plaque expansion speed and distance are comparable with the phage without LIN.

To follow the expansion of the infection front, we defined the following positions that characterize the infection propagation. (i)*r*_*PB*0_, defined as the largest *r*-value where the *R*(*r*_*PB*0_) = *B*_0_ (*P*(*r*_*PB*0_) = *B*_0_), where *B*_0_ is the initial bacterial concentration. *B*_0_ is the concentration of “microcolonies” if bacteria grow locally [10, 19, 20]. Hence, this is the distance at which there will be one phage per microcolony. This quantity characterises the furthest distance where infection is likely to happen. Note that when phage concentration is below this value, the infection event should become stochastic in reality, though our current model does not capture this stochasticity. (ii) *r*_*super*_, defined as the radius at which the local concentration of total concentration of bacteria is equal to that of the phage, *R*(*r*_*super*_) = *B*_*tot*_(*r*_*super*_) (*P*(*r*_*super*_) = *B*_*tot*_(*r*_*super*_)). Given the propagation profile, the secondary adsorption will be dominant for *r < r*_*super*_. Hence LIN is likely to be induced for the LIN-phage for *r < r*_*super*_. (iii) *r*_*half*_, defined as the point at which *B*_*tot*_(*r*_*half*_) = *B*_*tot*_(*r*_*max*_)*/*2, where *r*_*max*_ is the edge of the system where the phage has not reached. Hence, *r*_*half*_ is the estimate of the size of the visible zone of the killing. i.e., visible plaque size. (iv) *r*_*ZOI*_, defined as the point at which *B*(*r*_*ZOI*_) = *B*(*r*_*max*_)*/*2. This is similar to *r*_*half*_ but only looking at the uninfected cells. Hence, *r*_*ZOI*_ is the estimate of the size of the zone of infection.

The time evolution of *r*_*PB*0_, *r*_*super*_, *r*_*half*_, are *r*_*ZOI*_ are shown in Fig. 4 for *r*-mutant (C) and LIN-phage (D). We see that the *r*_*PB*0_, *r*_*half*_, *r*_*ZOI*_ curves are in parallel until the expansion slows down, i.e., all three positions expand at the same speed. *r*_*super*_ has a different slope because it depends both on the bacterial growth and the phage increase, but importantly, *r*_*super*_ is always clearly smaller than *r*_*PB*0_. This means a region with more bacteria than phage particles with not too low phage concentration always exists at the front of phage expansion, ensuring single infections without LIN at the expanding front. The plaque size (*r*_*half*_) and the zone of infection (*r*_*ZOI*_) reach the final value by 300 min. Comparing the *r*_*half*_ and *r*_*ZOI*_, we found that the visible plaque radius for the LIN-phage is 86 % of that of *r*-mutant, but the zone of infection radius is 96 % of that of *r*-mutant. Here, it should be noted that the variation of the total bacterial cell density *B*_*tot*_ over space for LIN-phage is considerably more graded than that of *r*-mutant (Fig. 4AB at *t* = 240min); the visible plaque size for the LIN-phage may be even smaller if significantly more than 50% drop of cell concentration is necessary to appear clear.

We tested how the final plaque size *r*_*half*_ and the zone of infection *r*_*ZOI*_ at time 300 min depends on LIN parameters, *f*_*τ*_ and *f*_*β*_. The values normalized to the same quantities for the *r*-mutant plaque are plotted in Fig. 5AB. For *f*_*τ*_ ≳ 3, virtually no cells lyse before nutrient depletion, and therefore, no detectable variation is observed for large *f*_*τ*_ regime. Interestingly, we find that the plaque radius of LIN-phage is about 85% of that of r-mutant while the zone of infection *r*_*ZOI*_ of LIN-phage is comparable with that of the *r*-mutant (ratio close to one) in most of the parameter space. The exception is when *f*_*τ*_ is close to 1 (almost no delay for LIN), which naturally makes the plaques of the LIN-phage bigger for larger *f*_*β*_. Since the unlysed LIN cells will not be able to grow but eventually lyse and produce a high number of phage particles, the comparable size of the zone of infection shows that LIN enables the phage to produce significantly more progeny without giving a disadvantage in infecting and killing bacteria cells in a spatially structured environment.

**Fig. 5:**
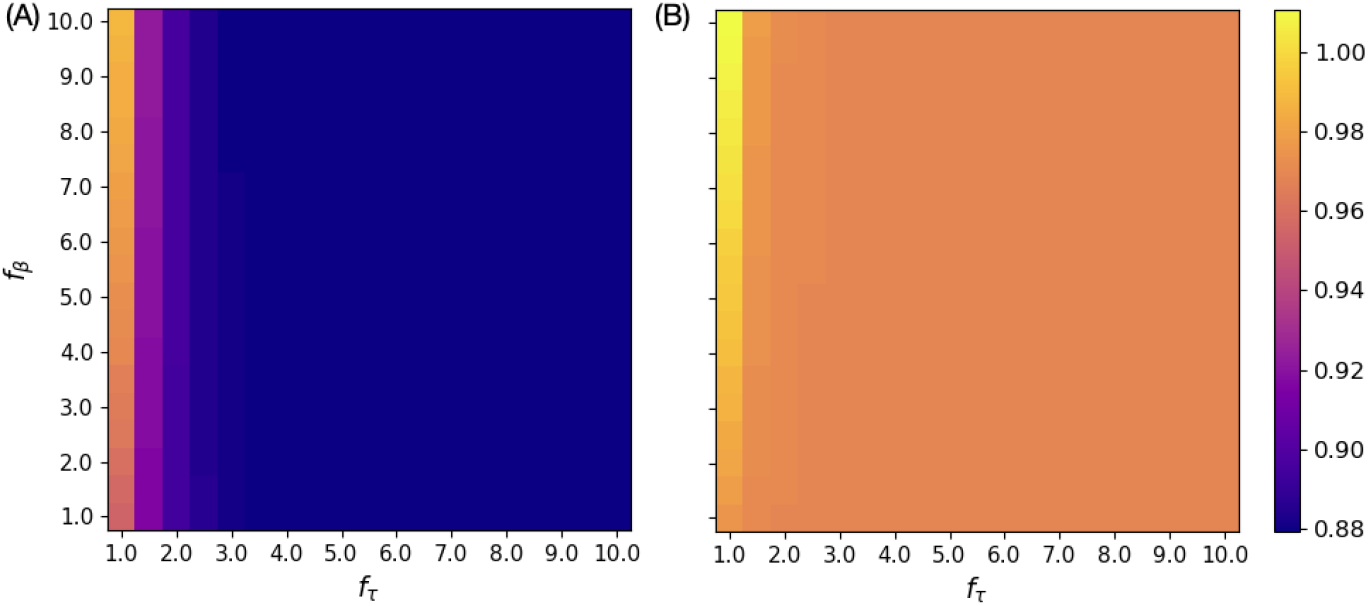
Plaque size and zone of infection dependence of lysis inhibition parameters. (A) Visible plaque radius *r*_*half*_ at 300 min normalized to that of the *r*-mutant. (B) Zone of infection *r*_*ZOI*_ at 300 min normalized to that of the *r*-mutant. The homogeneous color in *f*_*τ*_ ≳ 3 is because virtually no cells lyse before nutrient depletion.

## Discussion

We have constructed a simple mathematical model to simulate population dynamics of phage infection with LIN, where secondary adsorption extends latency time by a factor *f*_*τ*_ and burst size by factor *f*_*β*_. Simulation of the phage infection in an independent batch culture showed that the benefit (final phage number) of LIN is larger for smaller *f*_*τ*_ and larger *f*_*β*_, i.e., faster burst and larger burst size, as naturally expected.

However, in the competition between the *r*-mutant (non-LIN phage) and LIN-phage in the same batch culture, it was found that the longer *f*_*τ*_ is actually beneficial for the competition: LIN kicks in only after the system runs out of available uninfected bacteria cells, and after that, the secondary adsorption exclusion enables the LIN-state cells to inactivate the free *r*-mutant phages by adsorption, giving advantage for larger *f*_*τ*_ even without the burst size increase by LIN.

The LIN-phage are known to make visibly smaller plaques compared to the *r*- mutant. However, our simulation showed the zone of infection by LIN-phage reaches as far as that of *r*-mutant phages. The mechanism is simple: During the plaque expansion driven by the phage diffusion, the phage concentration drops towards the front while the bacterial density is higher in the uninfected region, and this structure makes secondary adsorptions unlikely at the front of infection. Hence, the burst is not delayed at the front, making the infection front unaffected by LIN. LIN is only induced behind the infection front where phage concentration is high enough, leaving the turbid region and making the apparent plaque, or the zone of cell lysis, smaller. The size of the zone of infection depends little on LIN parameters, demonstrating that this is a robust feature of the LIN-phage plaque.

This result is consistent with the interpretation that the plaque expansion in the current model is driven by the front of the expanding phage population as a pulled wave. It was suggested that when the diffusion of phages in a bacterial loan is severely reduced, the dynamics change to a pushed wave, where the growth of phage in the bulk of the wave becomes the driver of expansion [25]. In such a case, the plaque propagation of LIN-phage would be limited compared to the *r*-mutant since the phage population growth in the bulk is visibly lower (Fig. 4).

It should be noted that the final plaque appearance depends strongly on how the phage lysis and the burst size are reduced as the nutrient runs out. In the present simulation, we assumed that lysis stops when the nutrients run out, leaving the LIN-state cells unlysed. It is also possible to stop the plaque expansion upon starvation by making the burst size reach zero when the nutrients run out, leaving the latency time finite (cf. [26]). This may be closer to the reported bacterial growth rate dependence of the phage parameters for T4 by Hadas *et al*. [7], though their experiment was focused on the steady-state growth, while the growth rate changes continuously during the plaque expansion. In the current simulation, we chose to make the latency time diverge when the nutrient is unavailable so that the infected cells stay unlysed. We believe it is a reasonable assumption that most cells in the LIN state stay intact in a typical plaque assay to explain the smaller zone of clearance for the LIN-phage plaque than that of *r*-mutant. However, a detailed understanding of the bacterial physiology dependence of the latency time and the burst size both for single infected and superinfected cells is necessary for a more quantitative analysis of the plaque morphology. It will also help understanding dynamics of T4 attack on a growing colony [21].

Abedon [9] has studied the collision of T4 *r*-mutant plaques and wild type (WT)T4 (with LIN) plaques and reported the zone of interference, which appears that the *r*-mutant cannot destroy the cells too close to the WT T4 plaque centre. Deducing the radius of the interference, it has been proposed that WT T4 has infected almost as large a radius as the *r*-mutant but left them in an unlysed LIN-state. Our mathematical analysis qualitatively supports this hypothesis. Further detailed experimental research on T4 plaque structure will be beneficial to develop a more quantitative model for LIN phages.

We should also note that the analyzed model for LIN is quite simplified, where the secondary adsorption makes a fixed amount of delay in latency time *f*_*τ*_ and the change of burst size *f*_*β*_. In order to find a robust feature that is less dependent on the specific choice of the parameters, we focused on the features robustly observed for a large range of *f*_*τ*_ and *f*_*β*_. However, in reality, *f*_*τ*_ and *f*_*β*_ should positively correlate [5, 13]. Furthermore, *f*_*τ*_ depends on the extent of additional infection [3], and some literature suggests that every secondary adsorption delays the latency time for a certain duration, such that the timing and number of secondary adsorptions affect the latency time of LIN-cells [27]. In the liquid culture, the sudden clearance of LIN-cells after several hours, called LIN collapse, has also been observed, which could possibly be triggered by too many secondary adsorptions per cell [9]. We plan to establish a more detailed model for the LIN process and analyze the behaviour more quantitatively.

It is known that high multiplicity of infection (MOI) increases the frequency of the lysogenization upon infection of temperate phage *λ* [28–32]. It is natural to speculate that this is also because MOI signals epidemiological state [33, 34], in parallel to the currently analyzed secondary adsorption-dependent LIN [35, 36]. Our analysis indirectly suggests that MOI-dependent regulation of lysogenization is also a good strategy, even in a spatially structured environment, since infected cells keep lysing at the edge of the epidemic spreading.

Interestingly, *Bacillus subtilis* phages from the SP*β* family were found to alter their lysogenization frequency using a peptide-based communication system, termed arbitrium, produced upon infection, high levels of which promote lysogeny [37–39]. Because both arbitirum and MOI signal similar information in this scenario [35], we could ask why MOI was not enough; also, we could ask if there is an example in nature where LIN is regulated by an arbitirum-like system, and if not, why. It is tempting to think that they may carry somewhat different information. For example, their spatial resolution can be different; arbitrium should diffuse faster than phage particles due to their size, and therefore, arbitrium level could be influenced by infection in a larger area than where the infection is currently happening. It should be noted that arbitirum level also regulates the exit from lysogeny, where arbitirum is produced by prophages [40–42]; in this situation, arbitrium level signals lysogen concentration, which is information that MOI cannot provide. It is possible that the evolution of arbitrium may be driven by providing information that MOI cannot provide.

In conclusion, we proposed a simple mathematical model to simulate the spreading of LIN-phage among host bacteria. The simulation of the LIN-phage has shown that phage absorption by LIN-ed cells is a strong competitive advantage against an *r*-mutant, i.e. a phage that does not do LIN. We have also shown that the actual zone of infection by a LIN phage in plaque formation can be as far as the plaque size of *r*-mutant. The analysis suggests that secondary adsorption-triggered LIN is a robust strategy for the virulent phage both in well-mixed and structured environments.

## Data availability statement

The codes used for simulations and plotting are available at: https://github.com/UlrikHvid/Competitive-advantages-of-T-even-phage-lysis-inhibition-in-response-to-secondary-infection

## Acknowledgments

NM was supported by Novo Nordisk Foundation (NNF21OC0068775).

