## supplementary text for "Competitive advantages of T-even phage lysis inhibition in response to secondary infection"

**Supplementary Information: Competitive advantages of secondary adsorption-triggered lysis inhibition for T-even phages**

Ulrik Hvid and Namiko Mitarai

### S1 The model equations for plaque formation

Below we include the model equations for the plaque-formation models. The symbols used are in parallel with those for well-mixed systems in the main text, but the variables for plaque formation models are local concentration in two dimensions (i.e., per area), and we ignore the variation in the depth direction within the thin top agar of the depth $\Delta a$. In accordance with the unit conversion from volume concentrations to area concentrations, the phage adsorption constant $\eta$ is replaced with

$$\eta_s \equiv \eta/\Delta a \tag{S1}$$

In addition, the phage and nutrient equations include diffusion terms.

#### S1.1 The model equation for plaque formation of the $r$-mutant

For the $r$-mutant, the model equations (MP0) are given as follows.

$$\frac{\partial B}{\partial t} = g_n(n)\ B - \eta_s\ R\ B \tag{S2}$$

$$\frac{\partial I_1^R}{\partial t} = \eta_s\ R\ B - I_1^R \cdot N/\tau(n) \tag{S3}$$

$$\frac{\partial I_i^R}{\partial t} = (I_{i-1}^R - I_i^R) \cdot N/\tau(n),\ \ i \in [2, N-1] \tag{S4}$$

$$\frac{\partial I_N^R}{\partial t} = I_{N-1}^R \cdot N/\tau(n) - I_N^R \cdot N/\tau(n) \tag{S5}$$

$$\frac{\partial R}{\partial t} = \beta(n) I_N^R \cdot N/\tau(n) + (D_R \nabla^2 - \eta_s\ B_{tot})R \tag{S6}$$

$$\frac{\partial n}{\partial t} = -\frac{1}{Y} g_n(n)\ B + D_n \nabla^2 n \tag{S7}$$

$$\text{with } B_{tot} \equiv B + \sum_{i=1}^{N} I_i^R \tag{S8}$$

#### S1.2 The model equation for plaque formation of LIN-phage

For the LIN-phage, the model equations (MP1) are given as follows.

$$\frac{\partial B}{\partial t} = g_n(n)\ B - \eta_s\ P\ B \tag{S9}$$

$$\frac{\partial I_1^P}{\partial t} = \eta_s\ P\ B - \eta_s\ P\ I_1^P - I_1^P \cdot N/\tau(n) \tag{S10}$$

$$\frac{\partial I_i^P}{\partial t} = (I_{i-1}^P - I_i^P) \cdot N/\tau(n) - \eta_s\ P\ I_i^P,\ i \in [2, N-1] \tag{S11}$$

$$\frac{\partial I_N^P}{\partial t} = I_{N-1}^P \cdot N/\tau(n) - \eta_s\ P\ I_N^P - I_N^P \cdot N/\tau(n) \tag{S12}$$

$$\frac{\partial L_1}{\partial t} = \eta_s\ P\ \sum_{i=1}^N I_i - L_1 \cdot N/(f_\tau \tau(n)) \tag{S13}$$

$$\frac{\partial L_i}{\partial t} = (L_{i-1} - L_i) \cdot N/(f_\tau \tau(n)),\ i \in [2, N-1] \tag{S14}$$

$$\frac{\partial L_N}{\partial t} = L_{N-1} \cdot N/(f_\tau \tau(n)) - L_N \cdot N/(f_\tau \tau(n)) \tag{S15}$$

$$\begin{aligned} \frac{\partial P}{\partial t} = \beta(n) I_N^P \cdot N/\tau(n) + f_\beta \beta(n) L_N \cdot N/(f_\tau \tau(n)) \\ +(D_P \nabla^2 - \eta_s\ B_{tot}) P \end{aligned} \tag{S16}$$

$$\frac{\partial n}{\partial t} = -\frac{1}{Y} g_n(n)\ B + D_n \nabla^2 n \tag{S17}$$

$$\text{with } B_{tot} \equiv B + \sum_{i=1}^N (I_i^P + L_i) \tag{S18}$$

### S2 Competition of LIN-phage and $r$-mutant in mixed culture when $r$-mutant does not trigger LIN

Figure S1 shows the simulation results of competition between LIN-phage and $r$-mutant in liquid media when the secondary infection by $r$-mutant DOES NOT trigger LIN; LIN is triggered only when secondary adsorption of a LIN-phage happens to a bacterium which is first infected by a LIN-phage. The model equations are modified from that of MC1 by replacing $(R+P)$ in the secondary adsorption terms in eq. (25)-(28) with $P$. The rest of the simulation setup was the same as that for Fig. 2 in the main text.

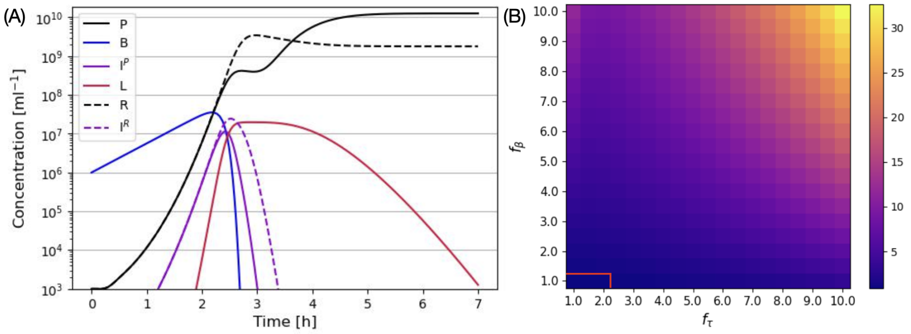


Figure S1: **Competition of $r$-mutant and LIN-phage in a mixed culture, when $r$-mutant cannot trigger LIN.** (A) An example time course of competition between $r$-mutant and LIN-phage with $f_\tau = f_\beta = 5$ in a mixed culture. (B) The final phage ratio between LIN-phage ($P$) and r-mutant ($R$) after 20 hours in a mixed culture. The red line shows the contour of the ratio being one. Note that the color bar is assigned differently from that of Fig. 2B, because the range of the fold difference is more extensive in this setup. LIN-phage is dominating in most of the parameter space.
